# Survival mechanism of a novel marine multistress-tolerant *Meyerozyma guilliermondii* GXDK6 under high NaCl stress as revealed by integrative omics analysis

**DOI:** 10.1101/2021.06.28.450280

**Authors:** Xinghua Cai, Huijie Sun, Huashan Bai, Yanyi Chen, Muhammad Kashif, Ru Bu, Xueyan Mo, Guijiao Su, Qian Ou, Bing Yan, Chengjian Jiang

## Abstract

A novel strain named *Meyerozyma guilliermondii* GXDK6 was provided in this work, which was confirmed to survive independently under high salt stress (12% NaCl) or co-stress condition of strong acid (pH 3.0) and high salts (10% NaCl) without sterilization. Its survival mechanism under high salt stress was revealed by integrated omics for the first time. Whole-genome analysis showed that 14 genes (e.g., *GPD1* and *FPS1*) of GXDK6 relevant to salt tolerance were annotated and known to belong to various salt-resistant mechanisms (e.g., regulation of cell signal transduction and glycerol metabolism controls). Transcriptome sequencing results indicated that 1220 genes (accounting for 10.15%) of GXDK6 were differentially transcribed (p < 0.05) when GXDK6 growth was under 10% stress for 16 h, including important novel salt-tolerant-related genes (e.g., *RTM1* and *YHB1*). Proteomics analysis demonstrated that 1005 proteins (accounting for 27.26%) of GXDK6 were differentially expressed (p < 0.05) when GXDK6 was stressed by 10% NaCl. Some of the differentially expressed proteins were defined as the novel salt-tolerant related proteins (e.g., sugar transporter STL1 and NADPH-dependent methylglyoxal reductase). Metabolomic analysis results showed that 63 types of metabolites (e.g., D-mannose, glycerol and inositol phosphate) of GXDK6 were up- or downregulated when stressed by 10% NaCl. Among them, D-mannose is one of the important metabolites that could enhance the salt-tolerance survival of GXDK6.

**IMPORTANCE:** Microbial contamination is a huge obstacle in industrial fermentation. The emergence of multistress-tolerant microorganism is expected to realize industrial fermentation without sterilization by controlling specific conditions. However, microorganisms eligible for non-sterile fermentation are required to survive independently under the selected special conditions for the fermentation conditions to be controlled to avoid microbial contamination. Here, a novel marine Meyerozyma guilliermondii was presented, which is able to survive well under high salt stress, its survival mechanism was systematically revealed by integrated omics technology. In addition, finding that NaCl stress could also stimulate the biosynthesis of functional metabolites from GXDK6 (e.g., calcitriol and didemnin B). Among the functional metabolites, calcitriol biosynthesis via microbial method was rarely reported. Thus, its biosynthetic mechanism was further revealed. The findings in this study contributed to understanding the survival mechanism of M. guilliermondii under high salt stress, and the development of new molecular drugs from M. guilliermondii GXDK6.

**Graphic abstract:** 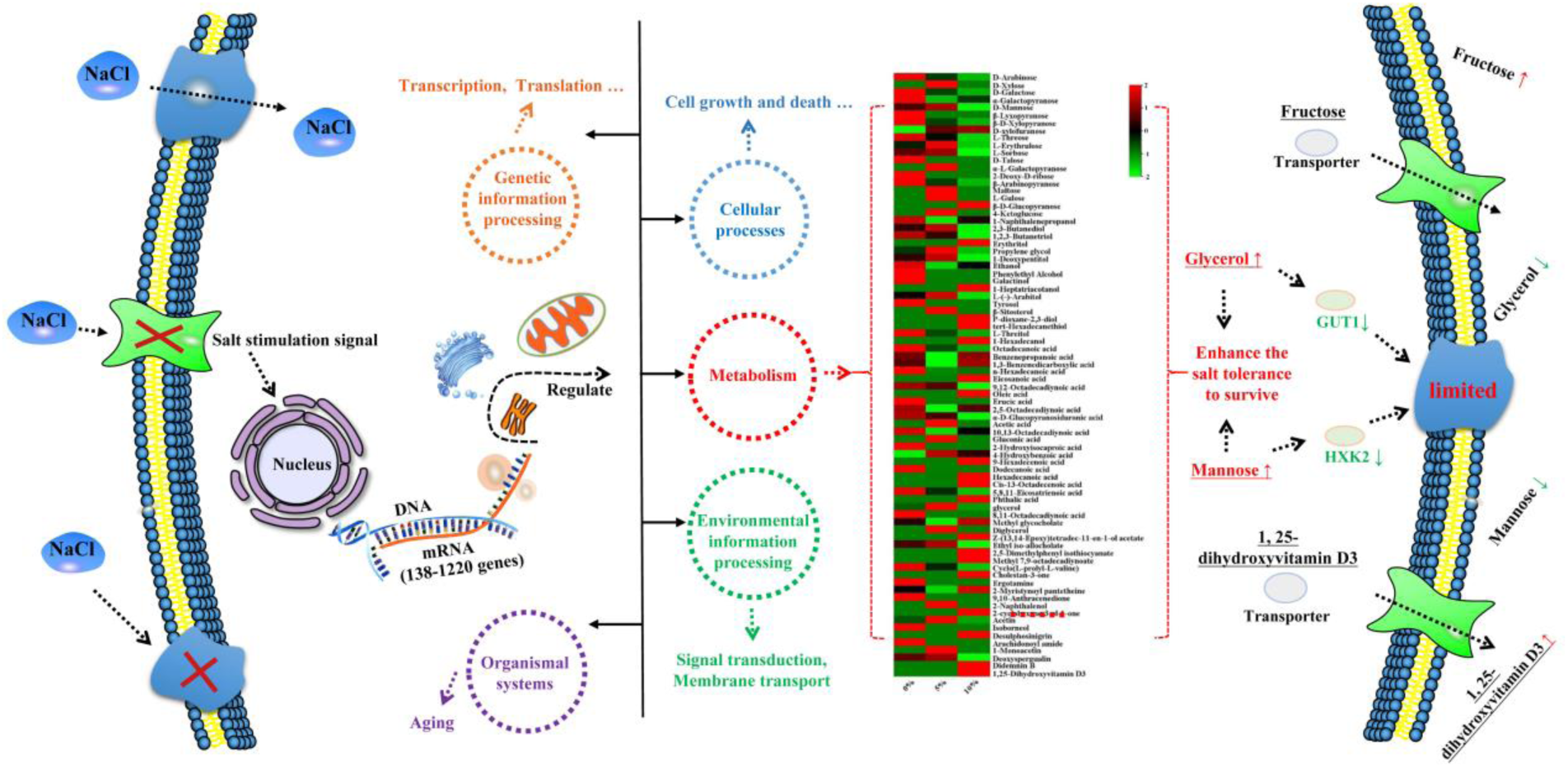

## BACKGROUND

Sterilization is important to avoid microbial contamination in the modern fermentation industry, which requires high facility investment and technical force (1). Given that non-sterile biological fermentation is achievable, it could greatly simplify the process and reduce production cost (2). However, microorganisms eligible for non-sterile fermentation are required to survive independently under the selected special conditions (e.g., strong acid, strong alkali and high salt) for the fermentation conditions to be controlled to avoid microbial contamination (3). Among them, adjusting the salt concentration in the fermentation medium may be one of the simplest and cheapest methods. In view of this, screening extremely salt-tolerant microorganisms (>10% NaCl) and revealing their salt-tolerance survival mechanism have gained more interests than before.

In response to salt stress, studies showed that microorganisms had evolved various strategies to withstand salt injury, including intracellular microenvironments, cell membranes, and metabolic levels (4–7). Yang et al. (8) presented a native salt-resistant *Meyerozyma guilliermondii* isolated from the mangrove ecosystem and further investigated its salt-resistance mechanism by comparative transcriptomics. Many key salt-tolerant genes were annotated and contributed to the survival of *M. guilliermondii* (e.g., FPS1 and GPD1). Similar results were also reported by Hou et al.(9), who revealed the salt tolerance mechanisms of *Zygosaccharomyces rouxiia* by comparative physiological and transcriptomic analyses and found that salt stress led to the accumulation of glycerol and trehalose and an increase in unsaturated fatty acid proportions. In addition, the genes involved in cellular metabolism and ribosome biosynthesis exhibited differential expression. However, few reports were found on the global effect of salt stress on the survival mechanism of *M. guilliermondii* (10). The relationship among the key genes, proteins, and metabolites relevant to salt tolerance remains unknown. Furthermore, previous reports hardly focused on the effect of functional secondary metabolites on the salt-tolerance survival of *M. guilliermondii* (11). Exploring how salt tolerance genes regulate the expression of salt tolerance-related proteins, what aspects of cell functions could be affected by these proteins, and whether it produces relevant metabolites conducive to the salt-tolerance survival of *M. guilliermondii* are urgent. The solutions to these bottlenecks could contribute to the survival mechanism of microorganisms under high salt stress (12, 13).

The previous study provided a novel multistress-tolerant *Meyerozyma guilliermondii* GXDK6, which showed broad pH tolerance (pH = 2.0–11.0) and high salt resistance (up to 14% NaCl or 18% KCl) (14). Its potential for non-sterile fermentation under high salt stress (12% NaCl) was also demonstrated. Due to its remarkable salt-tolerant survivability, it was hypothesized that GXDK6 may survive under high salt stress by regulating related genes to control the expression of salt-tolerant key proteins, which are beneficial to regulate cell function and/or produce some contributing secondary metabolites. Thus, to test this hypothesis, integrative omics strategies were performed to investigate the survival mechanism of *M. guilliermondii* GXDK6. This work contributes to understanding the metabolic regulation and helps develop some functional metabolites from *M. guilliermondii* GXDK6.

## RESULTS

### Salt tolerance and salt-removal ability of GXDK6

As shown in Fig. 1, GXDK6 was isolated from the subtropical marine mangrove sediments in the Beibu Gulf of China; its physicochemical characteristics were identified in previous studies (Figs. 1a–c) (14). The results showed that GXDK6 could continuously grow under a concentration of 0%–1% NaCl, indicating high salt tolerance (Fig. 1d). However, when the NaCl concentration was increased to 15%, the growth of GXDK6 was drastically inhibited, suggesting that the maximum tolerance of GXDK6 to NaCl may be up to be 15%. Thus, the salt-removal ability of GXDK6 was further investigated. As shown in Fig. 1e, the highest salt-removal rate of GXDK6 was 21.13%, but it decreased gradually with the increase in salt concentration. These findings indicated that GXDK6 has a potential application in the treatment of high-salinity wastewater (15). Fig. 1f shows the growth trend of GXDK6 under varying NaCl concentrations, demonstrating that high salt concentration substantially inhibits the growth of GXDK6, resulting in prolonged lag and logarithmic phases. In short, high salt stress changes the gene transcription expression of GXDK6 in the lag phase and probably affects cell metabolism in the logarithmic phase (5).

**Figure 1.**
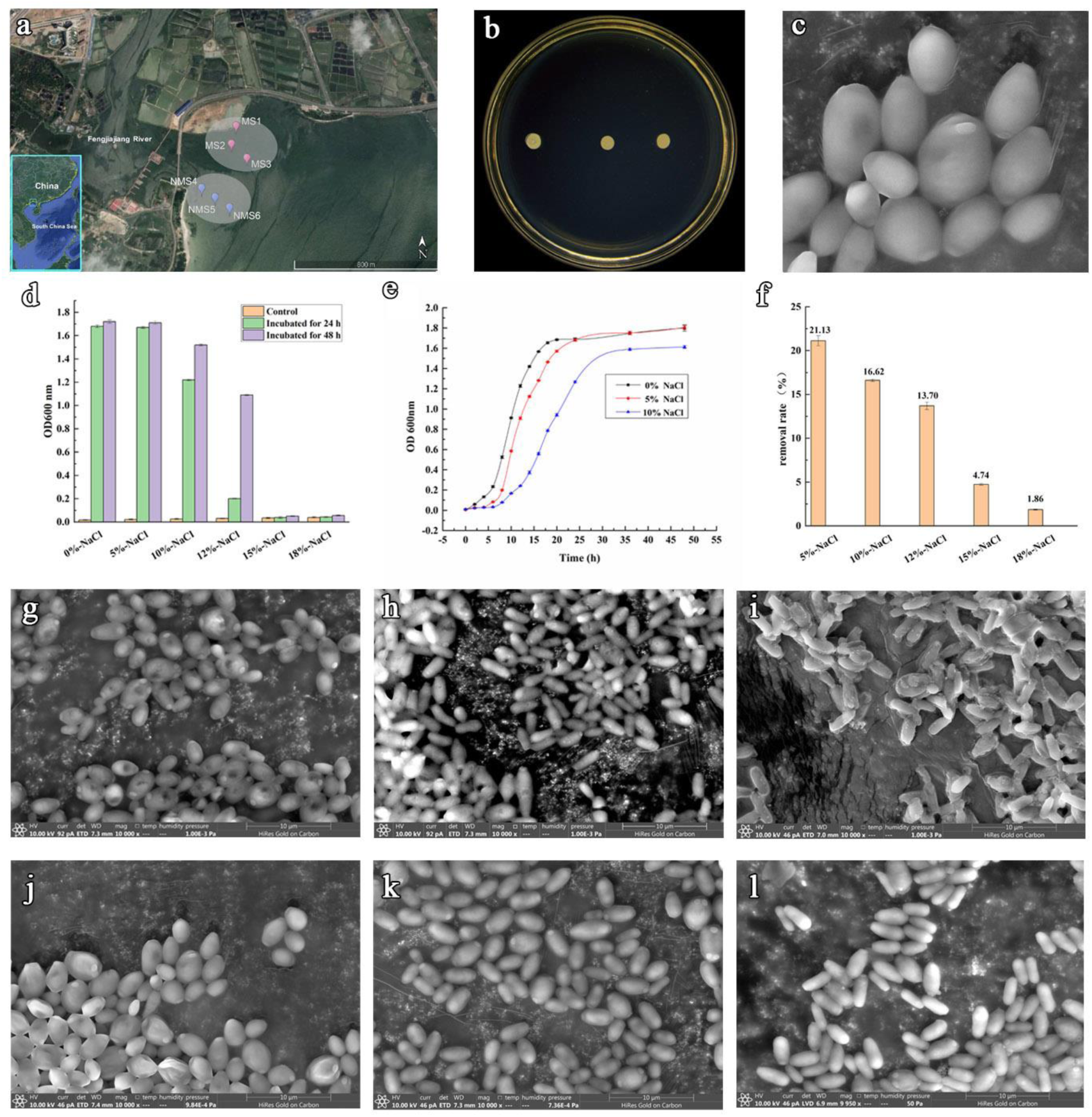
Physicochemical properties of GXDK6. (a) Sediments from the mangrove area in the Beibu Gulf of China; (b) colony morphology of GXDK6; (c) cell morphology of GXDK6; (d) salt tolerance of GXDK6; (e) salt-removal ability of GXDK6; (f) growth curve of GXDK6 under NaCl stress; (g) GXDK6 cells incubated for 16 h under 0% NaCl; (h) GXDK6 cells incubated for 16 h under 5% NaCl; (i) GXDK6 cells incubated for 16 h under 10% NaCl; (j) GXDK6 cells incubated for 48 h under 0% NaCl; (k) GXDK6 cells incubated for 48 h under 5% NaCl; and (l) GXDK6 cells incubated for 48 h under 10% NaCl.

### Surface morphology of GXDK6 under NaCl stress

The morphological changes in GXDK6 under NaCl stress were observed by scanning electron microscopy. The surface morphology of GXDK6 remained round or oval without NaCl stress (Figs. 1g and 1j). However, the cell morphology of GXDK6 gradually contracted and elongated when the NaCl concentration was increased to 10% (Figs. 1i and 1l), suggesting that the morphological changes in GXDK6 may respond to NaCl stress. Wang et al. (16) reported that lipid metabolism and membrane permeability are important in regulating the morphological changes in cells. This regulation is due to the signal instructions of gene transcription and expression, which allow cells to avoid internal damage effectively in high-permeability environments. However, how the detail regulatory signals work and whether this regulation leads to remarkable changes in metabolic patterns remain unknown.

### Genetic annotation of *M. guilliermondii* GXDK6

Whole-genome sequence analysis showed that the total bases of GXDK6 were 2,477,073,673 bps, and its GC content accounted for 38.88%. The genome sequencing data were deposited to the National Microbiology Data Center database (http://nmdc.cn) under accession number NMDC60014229 (14). Compared genomic analysis showed that 14 genes (Table 1) related to salt tolerance were annotated, suggesting that GXDK6 is a high salt-tolerant strain, well consistent with the results shown in Fig. 1.

**Table 1.**
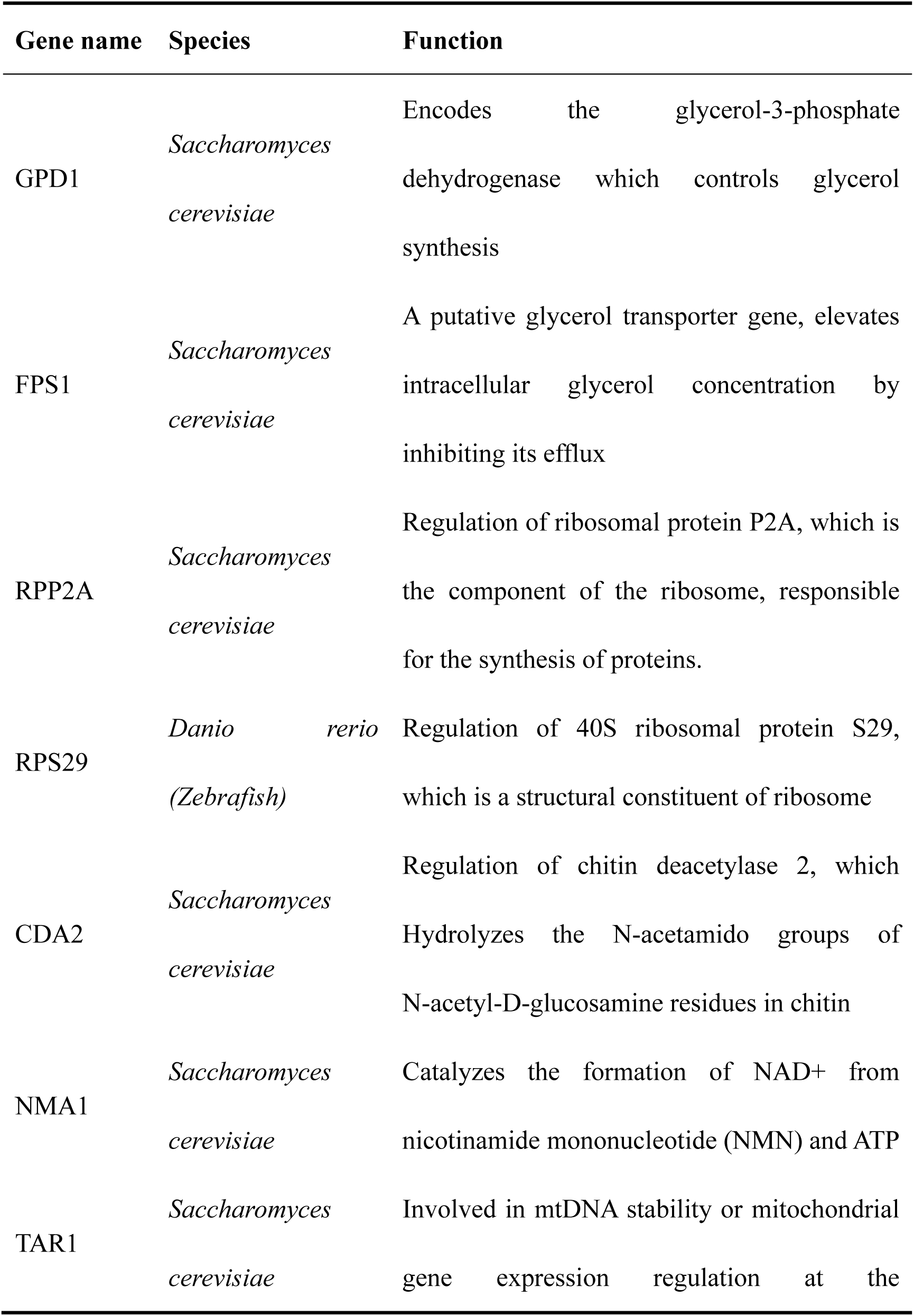
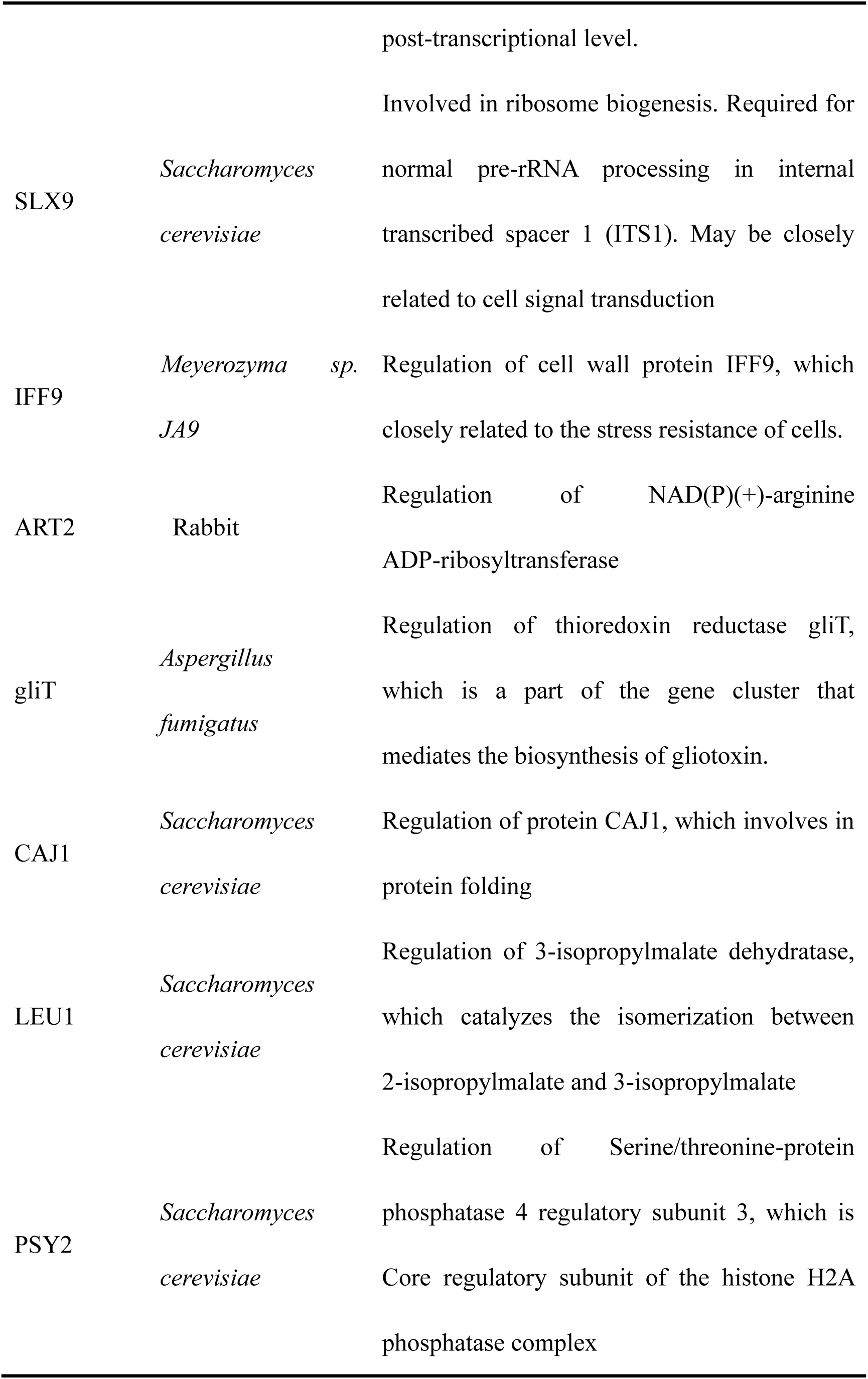
Annotation of the genes relevant to salt tolerance in *M. guilliermondii* GXDK6.

Among the salt-tolerant genes, *GDP1* and *FPS1* are considered the key genes of yeast response to salt stress. *GPD1* encodes glycerol-3-phosphate dehydrogenase, which controls glycerol synthesis (17). *FPS1*, a putative glycerol transporter gene, elevates intracellular glycerol concentration by inhibiting its efflux in yeast (18). These two key salt-tolerant genes play an important role in the adaptation to hyperosmotic stress dependent on signal transduction pathways, especially the transcription controlled by the yeast high-osmolarity glycerol response pathway, which mediates cellular adaptation to hyperosmotic stress and is responsible for glycerol production in yeast (19). In the case of hyperosmotic stress, yeast cells must react to the presence of external osmolytes that alter the osmotic pressure acting on the cell. A part of the response consists of the production of intracellular osmolyte glycerol, a compatible solute, to increase the internal osmolarity of the cell (20). In addition, glycerol plays vital physiological roles in the metabolism of the yeast, including osmoregulation and maintaining intracellular redox balance under an aerobic condition (21).

### Transcriptome sequencing and analysis of GXDK6 under NaCl stress

Transcriptome sequencing of GXDK6 was performed to reveal its gene regulation mechanism under NaCl stress. A total of 5175 reference genes were provided from the sequencing database, and the proportion of sequenced genes to reference genes were all higher than 99.07% (Table 2), indicating a high-quality transcriptome sequencing result (22), which provided a reliable basis for subsequent data analysis.

**Table 2.** Statistical analysis of the transcriptome sequencing genes.

| Samples | Refer<br>genes | Sequenced<br>refer genes (%) | Novel<br>Genes | Sequenced novel<br>genes (%) |
| --- | --- | --- | --- | --- |
| All | 5175 | 5172 (99.94%) | 44 | 44 (100.00%) |
| 0%-NaCl-16 h | 5175 | 5149 (99.50%) | 44 | 44 (100.00%) |
| 5%-NaCl-16 h | 5175 | 5127 (99.07%) | 44 | 44 (100.00%) |
| 10%-NaCl-16 h | 5175 | 5133 (99.19%) | 44 | 43 (97.73%) |
| 0%-NaCl-48 h | 5175 | 5167 (99.85%) | 44 | 44 (100.00%) |
| 5%-NaCl-48 h | 5175 | 5157 (99.65%) | 44 | 44 (100.00%) |
| 10%-NaCl-48 h | 5175 | 5153 (99.57%) | 44 | 44 (100.00%) |

As shown in Fig. 2, the transcription of 110 genes was upregulated and 459 genes were downregulated when GXDK6 was incubated in 5% NaCl for 16 h. Meanwhile, 622 genes were upregulated and 598 genes were downregulated when the NaCl concentration was increased to 10% (Fig. 2a). However, when the fermentation time was prolonged to 48 h, 226 genes were upregulated and 423 genes were downregulated when GXDK6 was incubated in 10% NaCl (Fig. 2c). Thus, the number of differentially expressed genes decreased with the prolongation of fermentation time, suggesting that GXDK6 takes time to regulate its gene expression in response to salt stress. Moreover, among the differential genes of GXDK6, 19–78 genes were common, regardless of whether they were incubated at 5% or 10% NaCl, (Figs. 2b and 2d). This finding suggested that these genes may participate in the same metabolic regulation process in GXDK6.

**Figure 2.**
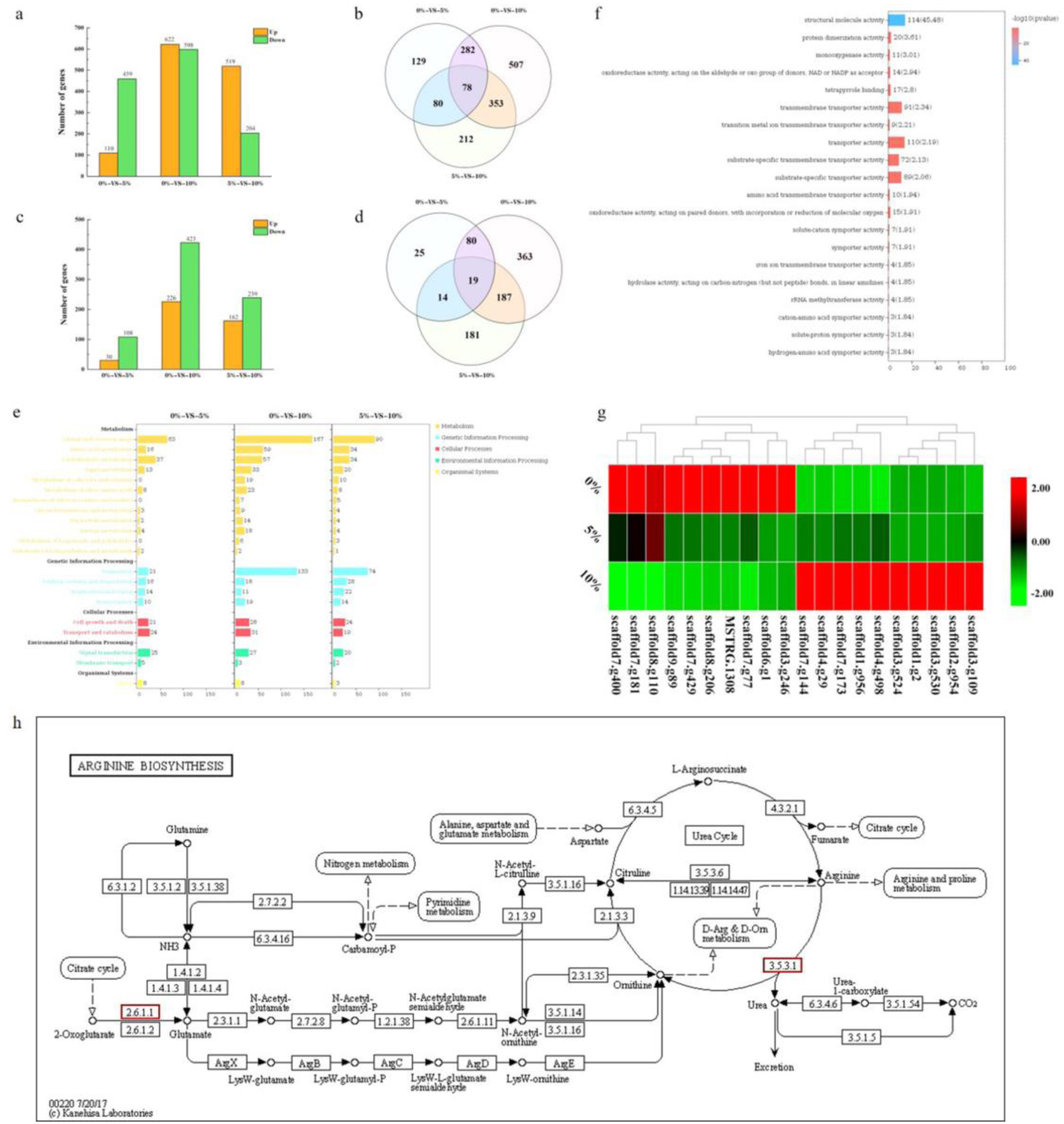
Transcriptome sequencing analysis of GXDK6. (a) Differential genes of GXDK6 under NaCl stress for 16 h; (b) Venn diagram analysis of the differential genes for 16 h; (c) differential genes of GXDK6 under NaCl stress for 48 h; (d) Venn diagram analysis of the differential genes for 48 h; (e) KO enrichment analysis of the differential genes for 16 h; (f) GO enrichment analysis of the differential genes for 16 h; and (g) regulatory pathway of *AAT2* (scaffold3.g530) in GXDK6.

KO enrichment analysis of the differentially transcribed genes were conducted, and the results showed that 229–525 genes, which were linked to five life activities (metabolism, genetic information processing, cellular processes, environmental information processing, and organismal systems) of GXDK6 were transcribed differentially by NaCl stress (Fig. 2e and Table 3). For example, when GXDK6 was stressed by 10% NaCl for 16 h, 247 genes (accounting for 4.77% of reference genes) relevant to cell metabolism were differentially transcribed. The most affected pathways were amino acid metabolism (accounting for 1.14% of reference genes), carbohydrate metabolism (accounting for 1.10% of reference genes), and lipid metabolism (accounting for 0.64% of differential genes). In terms of genetic information processing, the differentially transcribed genes were involved in regulating the translation (accounting for 2.57% of reference genes), folding, sorting and degradation (accounting for 0.35% of reference genes), replication and repair (accounting for 0.21% of reference genes), transcription (accounting for 0.37% of reference genes) of GXDK6. In cellular processes, 28 genes (accounting for 0.54% of reference genes) were transcribed differentially and involved in regulating the cell growth and death of GXDK6, while 31 genes (accounting for 0.60% of reference genes) were linked to its transport and catabolism. In terms of environmental information processing, 27 differential genes (accounting for 0.52% of reference genes) were relevant to the signal transduction of GXDK6, and three differential genes (accounting for 0.06% of reference genes) were involved in regulating the membrane transportation of GXDK6. Moreover, NaCl stress could lead to the differential transcription of eight genes (accounting for 0.15% of reference genes) relevant to the organismal systems of GXDK6, which are mainly involved in regulating the cell process of aging, indicating that salt stress could inhibit cell growth and/or cause cell apoptosis (23).

**Table 3.**
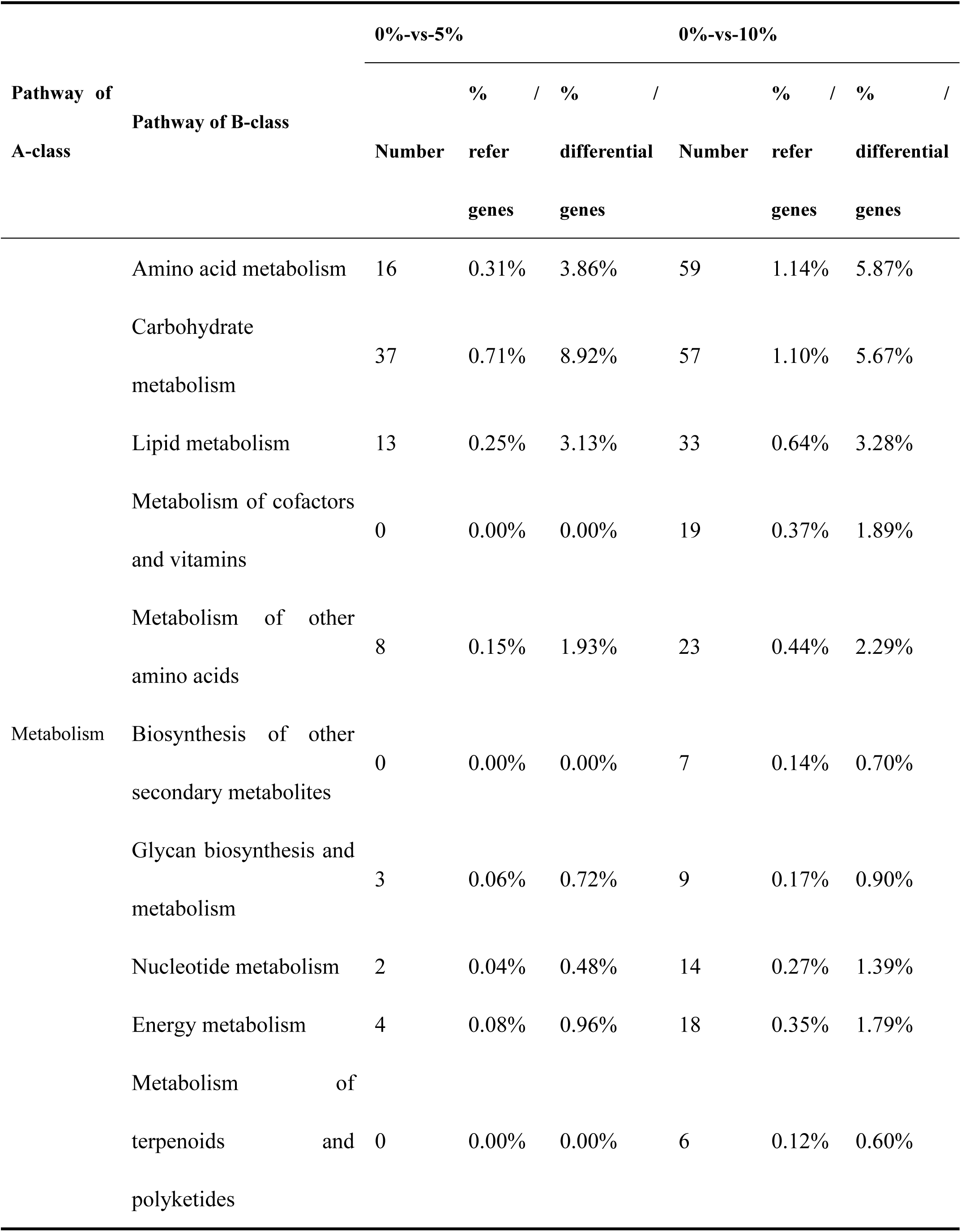
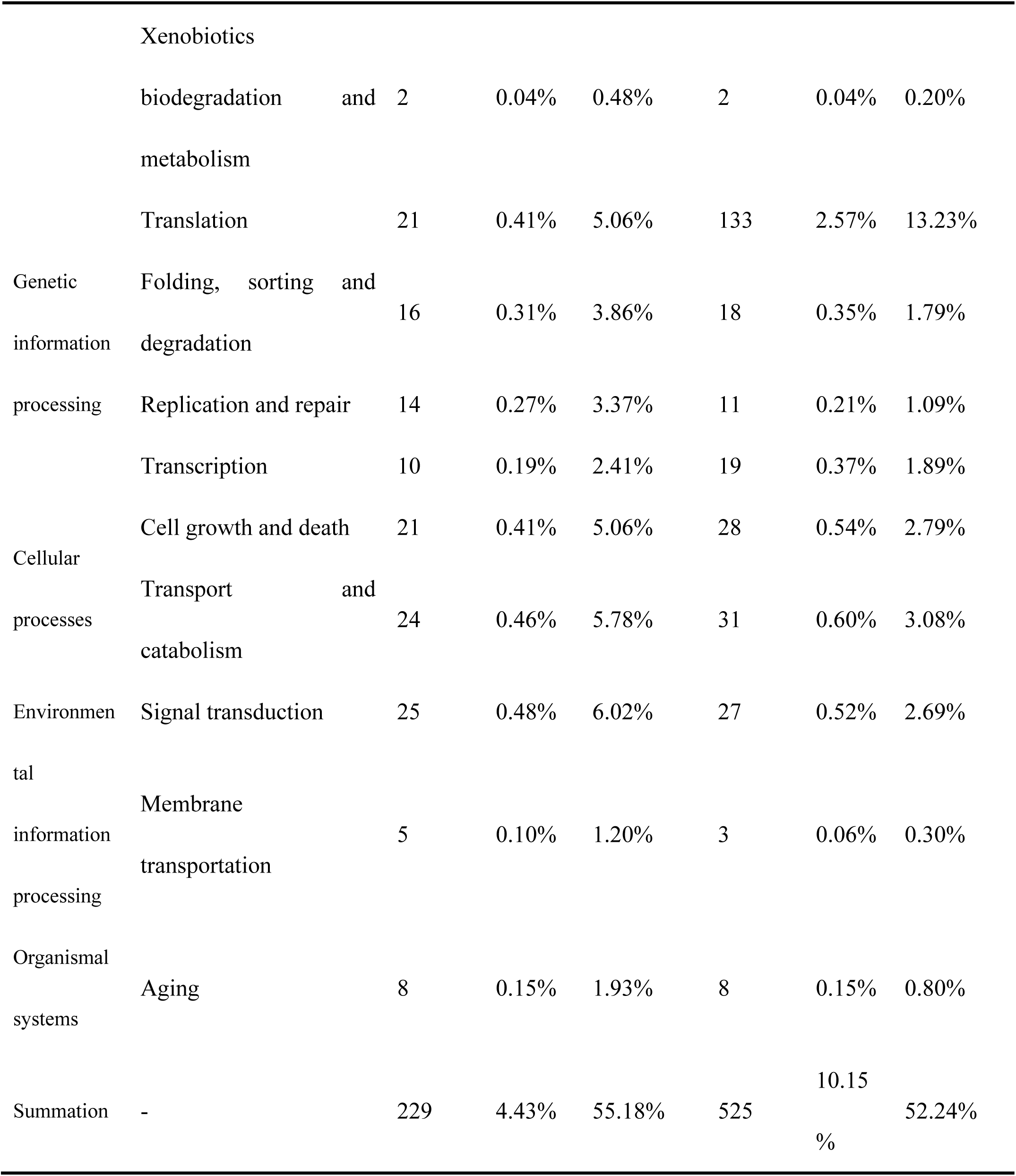
KO enrichment analysis of the differentially transcribed genes

GO enrichment analysis showed that the differential genes related to molecular function were mainly enriched in structural molecule activity (114 genes), transporter activity (110 genes), and transmembrane transporter activity (91 genes), as shown in Fig. 2f), indicating that the regulation of transport functional proteins is an important method for GXDK6 to survive in response to salt stress. The top 20 differential genes (Fig. 2g) were screened to further understand these regulatory genes, and the results showed that the highest upregulated gene was *RTM1* (scaffold4.g29, upregulated by 8.91-folds), which is involved in regulating the expression of protein *RTM1*. This protein confers resistance to GXDK6 to molasses, especially to the particular toxic element present in some molasses (24). The highest downregulated gene was *YHB1* (scaffold8.g110, downregulated by 14.95-folds), which is involved in regulating the expression of flavohemoprotein responsible for nitric oxide detoxification in an aerobic process. In the presence of oxygen and NADH, flavohemoprotein also showed NADH oxidase activity and contributed to regulating cell oxidative damage (25). Among the top 20 differential genes, *AAT2* (scaffold3.g530) was the most upregulated gene (4.54-folds) relevant to amino acid metabolism, which regulates the expression of aspartate aminotransferase to promote the biosynthesis of arginine (Fig. 2h) and contributes to the survival of GXDK6 under salt stress.

In summary, 5%–10% NaCl stress on GXDK6 could cause 4.43%–10.15% of genes to be differentially transcribed, indicating that the regulation of gene transcription (e.g., *RTM1*, *YHB1* and *AAT2*) and the transformation of cell metabolism (especially amino acid, carbohydrate, and lipid metabolisms) are important for GXDK6 to grow and metabolize in high-salinity environments (26). However, although transcriptome sequencing analysis of GXDK6 was performed, how these differentially transcribed genes regulate the expression of related proteins and whether they produce relevant metabolites conducive to the salt-tolerance survival of *M. guilliermondii* are still unknown. Studying the proteome and metabolome changes in GXDK6 under NaCl stress is urgent to explore these unsolved mysteries.

### Proteomics analysis of GXDK6 under NaCl stress

As shown in Fig. 3a, when GXDK6 was stressed by 5% NaCl for 16 h, 287 and 128 proteins were up- and downregulated respectively. The number of differentially expressed proteins (415) accounted for 11.51% of the total detected proteins (3604 proteins, provided in additional file 2). When NaCl concentration was increased to 10%, the number of differentially expressed proteins increased to 1005 (including 631 upregulated and 374 downregulated proteins), which accounted for 27.89% of the total detected proteins. Moreover, whether GXDK6 was stressed by 5% NaCl or 10% NaCl, 79 common proteins (accounting for 6.3%, Fig. 3b) were still found, suggesting that these differential proteins showed consistent resistance to salt stress. Subsequently, subcellular localization analysis of the differential proteins was further performed. The results demonstrated that when GXDK6 was grown under 10% NaCl, 37.5%, 25.1%, 18.4%, 7.9%, and 5.7% differential proteins were located in the nucleus, cytoplasm, mitochondria, plasma membrane, and extracellular matrix, respectively. These proteins accounted for 94.6% of the total detected proteins (Fig. 3c). This finding suggested that the nucleus, cytoplasm, and mitochondria are the main parts of GXDK6 in response to NaCl stress (the number of differential proteins located in these three parts accounted for 81% of the total detected proteins). Among them, nucleus may first receive the salt stimulation signals and then regulate the gene transcription (4.43%–10.15%) to control the biosynthesis and processing of proteins in ribosomes, the endoplasmic reticulum, and the mitochondria (27), thereby expressing several salt tolerance-related proteins (11.51%–27.89%), which remarkably contributed to the survival of GXDK6 under high salt stress.

**Figure 3.**
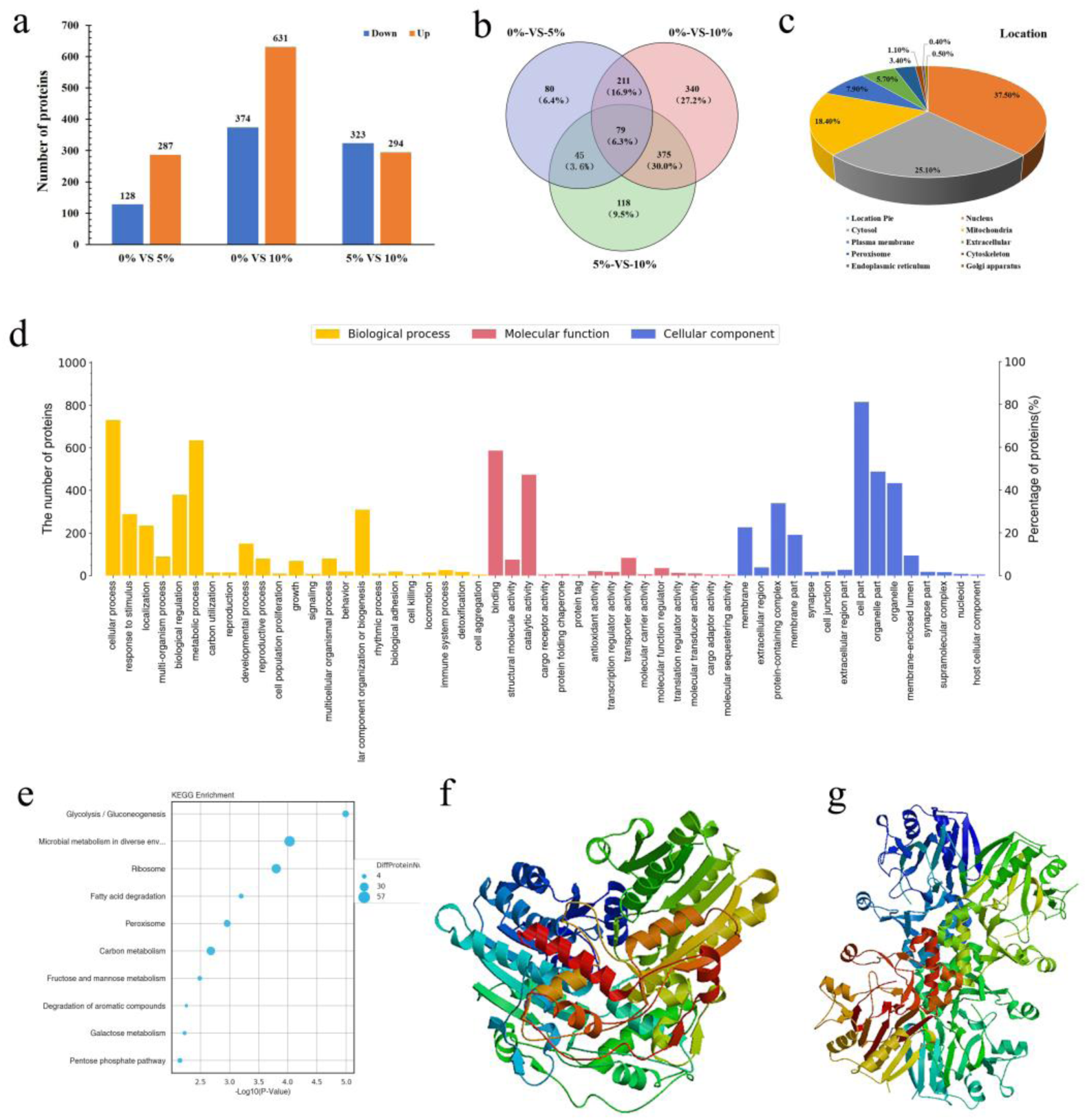
Proteomics analysis of GXDK6 under NaCl stress. (a) Differential proteins of GXDK6 under NaCl stress for 16 h; (b) Venn diagram analysis of the differential proteins; (c) subcellular localization analysis of the differential proteins; (d) GO enrichment analysis of the differential proteins; (e) KO enrichment analysis of the differential proteins; (f) sugar transporter STL1 (regulated by scaffold3.t827); and (g) NADPH-dependent methylglyoxal reductase (regulated by scaffold8.g110).

GO enrichment analysis was also performed in this work, and the results showed that the differentially expressed proteins of GXDK6 under NaCl stress were mainly enriched in the biological process (23 terms), molecular function (15 terms), and cellular component (15 terms, Fig. 3d). For example, when GXDK6 was grown under 10% NaCl stress for 16 h, the differential proteins were mainly enriched in the cellular process, metabolic process, protein binding, catalytic activity, and cell and organelle parts of GXDK6. The results of KO enrichment analysis (Fig. 3e) also showed that the differential proteins were mainly enriched in the microbial metabolism in diverse environments (57 proteins), glycolysis/gluconeogenesis (22 proteins), fatty acid degradation (11 proteins), and fructose and mannose metabolism (10 proteins). These findings suggested that the differential proteins mainly contributed to the cell structure regulation, substance transport, and metabolic pathway transformation of GXDK6, thus supporting GXDK6 to survive in high NaCl stress. They are also well consistent with the transcriptome sequencing analysis (Fig. 2) and results shown in Fig. 1.

The most upregulated protein, namely, sugar transporter STL1 (scaffold6.t443, upregulated by 1.73-folds, Fig. 3f), and the most downregulated NADPH-dependent methylglyoxal reductase (scaffold3.t827, downregulated by 2.15-folds, Fig. 3g) were further screened to obtain more details of the differential proteins. Sugar transporter STL1 is a sugar transporter responsible for balancing the sugar concentration and osmotic pressure of cell (28) and transporting the sugar that has accumulated too much or is not conducive to the salt-tolerance survival of GXDK6 (e.g., fructose). The function of NADPH-dependent methylglyoxal reductase is to catalyze the irreversible reduction of the cytotoxic compound methylglyoxal (MG, 2-oxopropanal) to (S)-lactaldehyde. MG is synthesized via a bypath of glycolysis and played a role in cell cycle regulation and stress adaptation (29). Therefore, when stressed by high salts, GXDK6 could quickly receive salt stress stimulus signals and then regulate its gene transcription (4.43%–10.15%) to control the expression of relevant proteins (11.51%–27.89%), thereby regulating its biological process, molecular functions, cell components, and metabolic pathways to maintain good growth under high salt stress. However, what changes in the metabolic pathways could contribute to the salt-tolerance survival of GXDK6 and whether some metabolites produced could enhance the survival of GXDK6 under high salt stress must be explored. Thus, metabolomics analysis of GXDK6 under NaCl stress was performed to solve these problems.

### Metabolomics analysis of GXDK6 under NaCl stress

As shown in Fig. 4, when GXDK6 was incubated in 0%–10% NaCl the types of its metabolic products considerably changed (provided in Additional file 3: Table 1S and 2S). The metabolites were classified into six categories, namely, saccharides, alcohols, phenols, organic acids, lipids, and other organic compounds (Fig. 4a). A total of 42, 45, and 45 fermentation products were found when it was grown under 0%, 5%, and 10% NaCl for 16 h, respectively, whereas 68, 56, and 55 fermentation products were identified when the incubation time was prolonged to 48 h, respectively (Fig. 4b). Among them, glycerol, phenol, and 2,4-di-tert-butylphenol were found as the common metabolites, suggesting that regardless of whether GXDK6 was stressed under 0%–10% NaCl, it still shared the same metabolic pathways.

**Figure 4.**
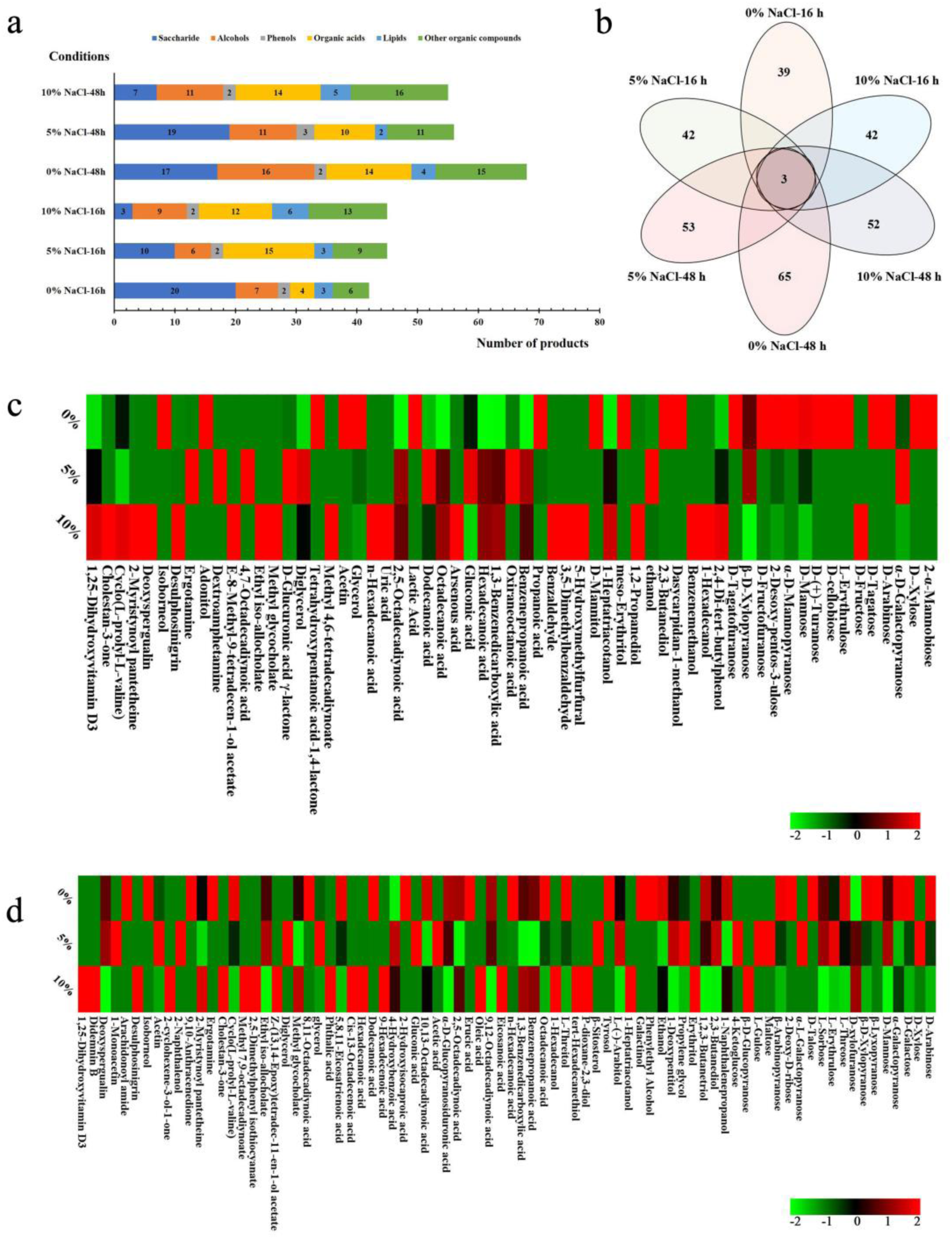
Metabolomics analysis of GXDK6 under NaCl stress. (a) Classification analysis of the metabolites; (b) total detected metabolites of GXDK6 under NaCl stress; (c) heatmap analysis of differential metabolites under NaCl stress for 16 h; and (d) heatmap analysis of differential metabolites under NaCl stress for 48 h.

Heatmap analysis of the differential metabolites is presented in Figs. 4c and 4d. When GXDK6 was stressed by 5%–10% NaCl for 16 h, 63 types of products were found to be considerably upregulated or downregulated. For example, 20 carbohydrates were detected when GXDK6 was incubated without NaCl stress. However, the number of carbohydrates decreased to 10 and three types with the increase in the NaCl concentration to 5% and 10%, respectively (Fig. 4a), indicating that the carbohydrate metabolism of GXDK6 was remarkable weakened or the metabolic pathways were transformed by NaCl stress. In addition, the types of organic acids (12) and lipids (six) increased by more than 2-folds when GXDK6 was stressed by 10% NaCl. Therefore, the accumulation of organic acids and lipids is likely beneficial to GXDK6 to increase its resistance to salt stress (30). Among the metabolites, glycerol was found as the consistent product when the NaCl concentration was increased from 0% to 10%; the corresponding glycerol decreased from 14.99% to 8.88%, with a decrease by 40.76%, demonstrating that glycerol is one of the important substances in response to salt tolerance (31). In addition, adding exogenous glycerol could enhance the salt-tolerance survival of GXDK6 (Additional file 3: Fig. 5S).

**Figure 5.**
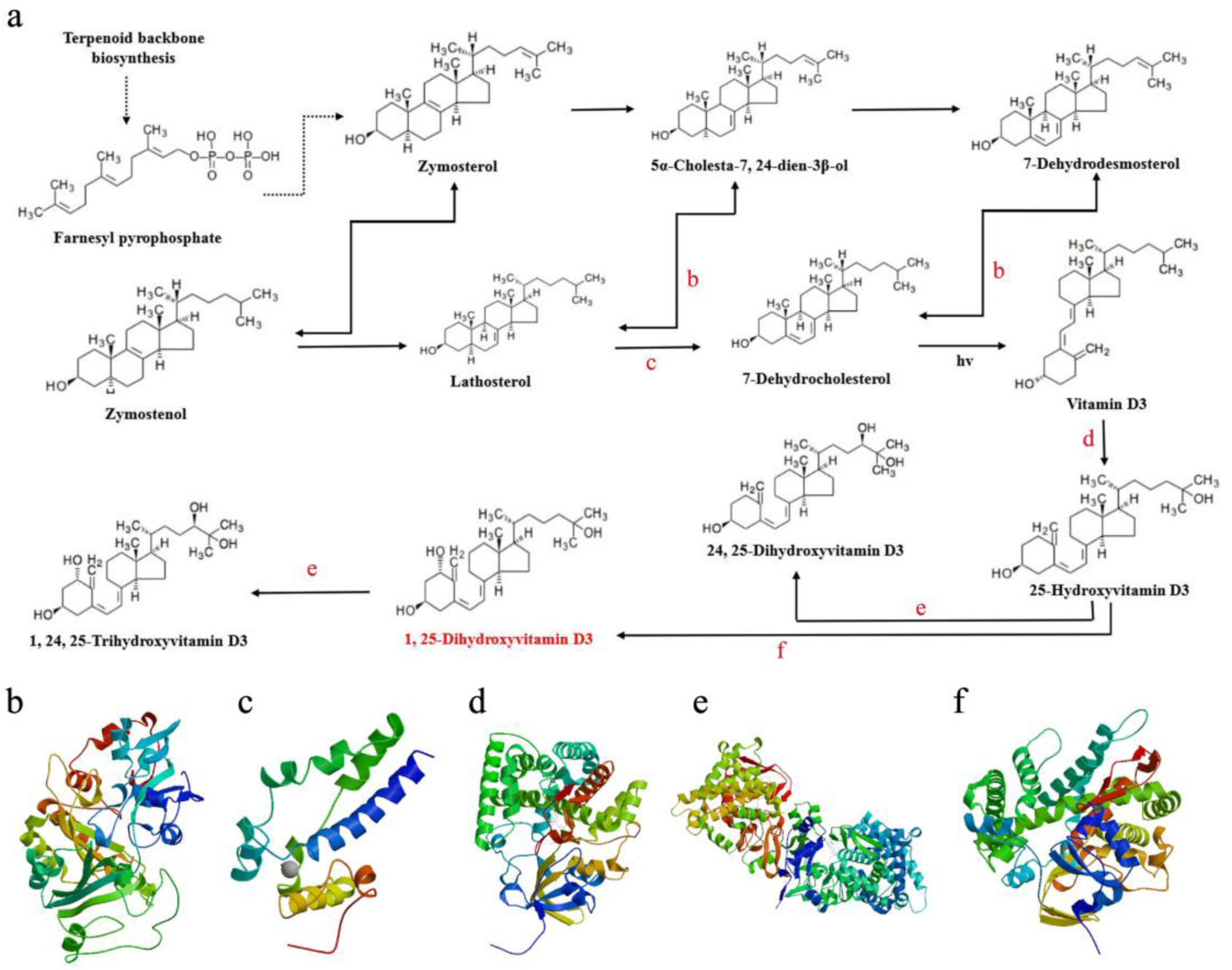
Biosynthesis mechanism of 1,25-dihydroxyvitamin D3. Metabolic pathway of (a) 1,25-dihydroxyvitamin D3, (b) delta (24)-sterol reductase, (c) C-5 sterol desaturase, (d) vitamin D3 dihydroxylase, (e) 1,25-dihydroxyvitamin D3-24-hydroxylase, and (f) 25-hydroxyvitamin D-1 alpha hydroxylase.

Moreover, the fermentation products of GXDK6 significantly changed under 5% g–10% NaCl stress compared with the control (p < 0.01), suggesting that these altered metabolites are beneficial to increasing the salt tolerance of GXDK6. For example, when GXDK6 was stressed by 10% NaCl for 16 h, the main upregulated metabolites were D-fructose, 2,4-di-tert-butylphenol, 1,25-dihydroxyvitamin D3; while the main downregulated metabolites were D-mannose, glycerol, and propanoic acid. These metabolites were regarded as important contributors to the salt resistance survival of GXDK6. The above transcriptome and proteome analysis results showed that D-fructose and D-mannose metabolisms were affected by the upregulated *YALI0B16192g*. Fructose was synthesized intracellularly, and it could be excreted outside the cell by fructose transporter (which could be encoded by *GLUT5* and *BmST1* and was not differentially expressed by NaCl stress). However, although mannose could also be synthesized intracellularly, the regulatory gene *HXK2* responsible for the transport of mannose was significantly downregulated (1.06-folds, p < 0.05), resulting in the weakening of the extracellular transport of mannose and significant downregulation of the extracellular mannose concentration (p < 0.05). In the metabolic pathway of fructose and mannose metabolisms, five key enzymes, namely, hexokinase (encoded by *HXK2*, which was downregulated by 1.06-folds), aldehyde reductase, L-iditol 2-dehydrogenase, D-iditol 2-dehydrogenase, and sorbose reductase (encoded by *YALI0B16192g*, which was upregulated by 3.74-folds), were involved in the regulation process. Furthermore, D-mannose was a vital biomarker that participated in regulating the salt tolerance of GXDK6. Adding exogenous 0.2% D-mannose could enhance the salt-tolerance survival of GXDK6 (Additional file 3: Fig. 6S).

**Figure 6.**
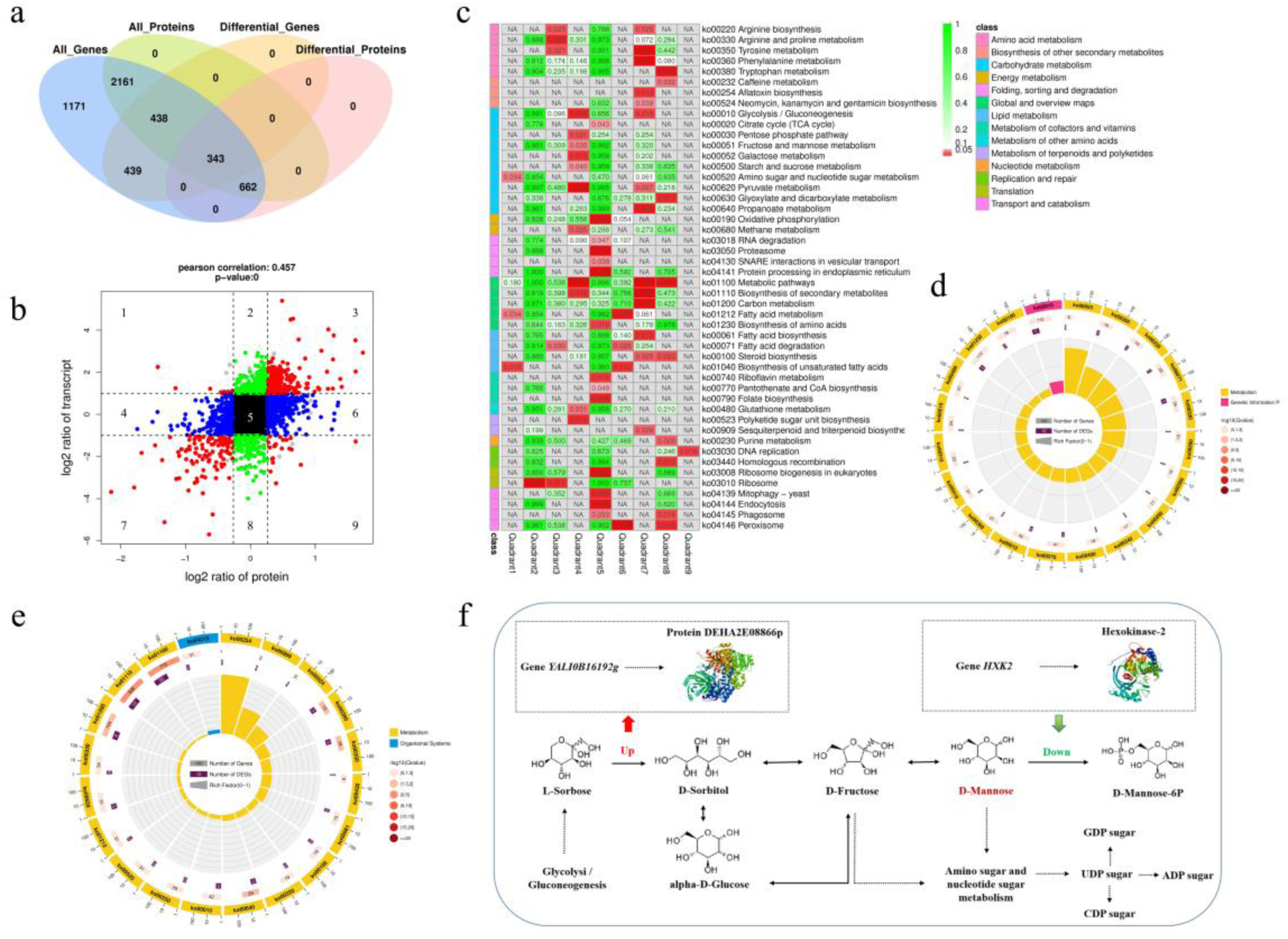
Association analysis of multi-omics data. (a) Association analysis of genes and proteins; (b) nine-quadrant analysis of differential genes and proteins; (c) KO enrichment analysis of differential genes and proteins in different quadrants; (d) KO enrichment analysis differential genes and proteins in the third quadrant; (e) KO enrichment analysis differential genes and proteins in the seventh quadrant; and (f) fructose and mannose metabolism regulated by differential genes and proteins.

Furthermore, a novel finding is that NaCl stress could stimulate GXDK6 to synthesize some important metabolites, such as calcitriol and didemnin B, which have been reported as potential valuable pharmaceutical ingredients (32, 33). Few reports also showed that these metabolites could be biosynthesized via microbial method (34, 35). Among them, calcitriol played an important role in cell proliferation and immune system, and its biosynthetic process was further revealed in this work (Fig. 5). The results showed that the upstream source of calcitriol was vitamin D3 (36), which was catalyzed by vitamin D3 dihydroxylase and then by 25-hydroxyvitamin D-1 alpha hydroxylase to synthesize calcitriol. Furthermore, calcitriol could be further catalyzed by 1,25-dihydroxyvitamin D3 24-hydroxylase to synthesize 1,24,25-trihydroxyvitamin D3. Six key enzymes, namely, C-8 sterol isomerase (coded by *ERG2*), delta (24)-sterol reductase (coded by *D6D02*), C-5 sterol desaturase (coded by *PAS_chr1*), vitamin D3 dihydroxylase (coded by *cyp105A1*), 1,25-dihydroxyvitamin D3 24-hydroxylase (coded by *Cyp24a1*), and 25-hydroxyvitamin D-1 alpha hydroxylase (coded by *B0E38*), were involved in the synthesis and metabolism of calcitriol. Therefore, this work also provided a new synthesis method for calcitriol production by microbial strategy under NaCl stress.

### Association analysis of the multi-omics data

As shown in Fig. 6, association analysis of the multi-omics data was performed, and the results showed that 3604 genes (69.12%) and proteins (100%) of GXDK6 were associated, suggesting a high-quality correlation analysis of the omics data (Fig. 6a). They were divided into nine quadrants to investigate how these associated genes and proteins work (Fig. 6b). The first, second, and fourth quadrants indicated that the expression abundance of proteins were lower than that of mRNAs, suggesting that they belonged to post-transcriptional or translational level regulation, such as miRNA regulation of targeted genes to inhibit the translation of related proteins. The third (upregulated) and seventh quadrants (downregulated) indicated the same trend of differential expression of mRNAs and proteins. The fifth quadrant indicated that neither mRNAs nor proteins were differentially expressed. The sixth, eighth, and ninth quadrants indicated that the expression abundance of proteins was higher than that of mRNAs, suggesting that they were a process of post-transcriptional or translational level regulation or accumulation of proteins. Therefore, the third and seventh quadrants were chosen as the main analysis objects in this work.

KO enrichment analysis of the associated genes and proteins from the nine quadrants was subsequently conducted (Fig. 6c). The results showed that the top five enrichment pathways were amino acid metabolism (especially in ko00220, ko00330, and ko00350), biosynthesis of other secondary metabolites (especially in ko01110, ko00100, and 01040), carbohydrate metabolism (especially in ko00010, ko00020, and ko00051), lipid metabolism (especially in ko00061, ko00071, and ko01212), and energy metabolism (especially in ko00190 and ko00680). Among the nine quadrants, the third quadrant showed that the upregulated genes and proteins were significantly enriched in the pathway of ascorbate and aldarate metabolism (ko00053, Fig. 6d), while the downregulated genes and proteins of the seventh quadrant were significantly enriched in the pathway of aflatoxin biosynthesis (ko00254, Fig. 6e). However, three common pathways, namely, arginine biosynthesis (ko00220), tyrosine metabolism (ko00350), and phenylalanine metabolism (ko00360), were found in the third and seventh quadrants, suggesting that the metabolic flux conversion of arginine, tyrosine, and phenylalanine are also important factors to promote the survival of GXDK6 under salt stress.

Moreover, the data from transcriptomic, proteomic and metabolomic analysis were combined, and the results showed that fructose and mannose metabolism is a typical pathway of GXDK6 in response to salt stress. GXDK6 upregulated the expression of *YALI0B16192g* and DEHA2E08866p and downregulated that of *HXK2* and hexokinase-2 to synthesize abundant fructose and mannose (Fig. 6f). D-fructose was further metabolized and transported to the outside of the cell, while D-mannose accumulated in a large amount in the cell due to the downregulation of the relevant gene and protein, thus contributing to the enhancement of the survivability of GXDK6. This result was further confirmed by the exogenous addition of D-mannose (Additional file: Fig. 6S).

## DISCUSSION

Microbial contamination is an obstacle to fermentation production of foods and molecular drugs. Current practices, such as process sterilization or antibiotic dosage, carry excess costs or encourage the development of antibiotic resistance. Developing microorganisms eligible for non-sterile fermentation shows a great application value by simplifying the process and reducing production cost (2). Although the currently known reports indicated that non-sterile fermentation is achievable (37), the selected microorganisms often could not be independently fermented with the corresponding culture conditions. Miscellaneous bacteria more or less still share the culture medium and exist in some uncertain biological fermentations, and this is not allowed for food and medical fermentation. Therefore, here, a probiotic identified as *M. guilliermondii* GXDK6 was presented. The findings confirmed that it could survive independently under the condition of 12% NaCl stress or co-stress condition of strong acid (pH 3.0) and high salt (10% NaCl) without sterilization. Only adjusting the salt concentration to avoid microbial contamination is considered one of the convenient methods. Thus, revealing the salt-tolerance survival mechanism of *M. guilliermondii* GXDK6 is beneficial to precisely control its salt-tolerant fermentation.

Most of the previous studies on *M. guilliermondii*, were focused only on the salt-tolerant-related gene (6–8), less on the relationship among the key genes, proteins, and metabolites (38), the effect of functional secondary metabolites on the salt-tolerance survival of *M. guilliermondii* were rarly reported. Therefore, a novel multistress-tolerant *M. guilliermondii* GXDK6, which was identified from subtropical mangrove sediments, was presented in the present study. Its survival mechanism under high NaCl stress was further revealed by integrative omics strategies (genomics, transcriptomics, proteomics, and metabolomics).

In this work, whole-genome sequencing analysis of *M. guilliermondii* GXDK6 was conducted, and salt-tolerant genes (including *GPD1, FPS1, CDA2,* and *IFF9*) were targeted. The annotated salt-tolerant genes were known to belong to various regulation mechanisms, including increasing the intracellular concentration of glycerol by inhibiting its efflux, regulating cell signal transduction involved in mtDNA stability or mitochondrial gene expression regulation at the post-transcriptional level, regulating the biosynthesis of chitinase, and enhancing the stress resistance of cells. These salt-tolerant genes and their regulatory mechanism were less demonstrated in a previous study; however, a part of the salt-tolerance genes in GXDK6 was reported, which still provided genetic basis for further understanding of the salt-tolerance mechanism of GXDK6 (39).

On the basis of the whole-genome analysis, the gene transcription differences in GXDK6 at 0%–10% NaCl stress were further explored. Numerous differentially transcribed genes (1220 genes, accounted for 23.57% of reference genes) were discovered when GXDK6 was stressed by 10% NaCl. These differential genes (except for genes with unknown functions) were mainly involved in regulating the metabolism (20.48%), genetic information processing (14.70%), cellular processes (10.84%), environmental information processing (7.22%), and organismal systems (1.93%) of GXDK6. Through a pathway enrichment analysis of the differential genes, their metabolic regulatory network was further revealed, and the contribution of the differential genes to the survival mechanism of GXDK6 was proven. Many novel salt-tolerant-related genes (e.g., *RTM1* and *YHB1*) were identified in GXDK6, thus further enhancing the understanding of salt tolerance-related genes (40).

In terms of proteomics analysis, 415–1005 proteins (accounted for 11.51%–27.89% of total detected proteins) relevant to the salt-tolerance mechanism of GXDK6 were identified, including the novel key salt-tolerant-related proteins, such as sugar transporter STL1 and NADPH-dependent methylglyoxal reductase. These differential proteins mainly contributed to the cell structure regulation, substance transport, and metabolic pathway transformation of GXDK6, thus supporting GXDK6 to survive in high NaCl stress. Moreover, an association analysis of the transcriptome and proteome data was conducted, and the results showed that 3604 genes (accounted for 69.64% of reference genes) and 3604 proteins (accounted for 100% of total detected proteins) were associated, suggesting that approximately 30.36% of genes may not be responsible for regulating the expression of proteins. The differentially expressed proteins mainly influenced the pathway of the amino acid metabolism, lipid metabolism, carbohydrate metabolism, and fatty acid metabolism of GXDK6.

At the metabolome level, 63–81 types of metabolites of GXDK6 were remarkably changed by NaCl stress. These metabolites were mainly produced from the carbohydrate metabolism, amino acid metabolism, and lipid metabolism of GXDK6. Glycerol, mannose, and calcitriol were shown to be closely related to cell osmotic pressure regulation, stress resistance, and immune regulation, respectively, indicating their important contribution to the salt-tolerance survival of GXDK6 (41). GXDK6 was incubated in high NaCl stress containing exogenous mannose or glycerol to further reveal the function of important metabolites, and the final results showed that adding 0.2% mannose or 120 mg/L glycerol is beneficial to enhance the survivability of GXDK6 under high NaCl stress.

In summary, GXDK6 could quickly receive the salt stress stimulus signals under 5%–10% NaCl stress and then regulate its gene transcription (4.43%–10.15%, e.g., *RTM1, YHB1*) to control the expression of relevant proteins (11.51%–27.89%, e.g., sugar transporter STL1 and NADPH-dependent methylglyoxal reductase). These differential proteins contribute to adjust the biological process, molecular functions, cell components, metabolic pathways (especially the amino acid metabolism, carbohydrate metabolism, and lipid metabolism) and produce many important salt-tolerant-related metabolites (e.g., D-mannose and glycerol) that are helpful to the survival of GXDK6 under high salt stress. In addition, the transformation of metabolic flow under NaCl stress stimulates GXDK6 to biosynthesize various important molecular drugs (e.g., calcitriol and ascidianin B). This novel discovery could be beneficial for constructing new synthetic methods for these molecular drugs, and it shows great value and significance.

## MATERIALS AND METHODS

### Species and reagents

*M. guilliermondii* GXDK6 was identified from the subtropical mangrove sediments in the Beibu Gulf of the South China Sea (21°29′25.74″ N, 109°45′49.43″ E). It is now deposited in the China General Microbiological Culture Collection Center (CGMCC) under CGMCC no. 16007 (14). N-methyl-trimethyl-silyl-trifluoroacetamide (MSTFA) and methoxyamine pyridine hydrochloride were of chromatographic grade and purchased from Sigma–Aldrich, Inc. (Darmstadt, Germany). Glucose, yeast powder, agar powder, peptone, HCl, NaOH, and NaCl were of analytical grade and purchased from Novagen (Darmstadt, Germany).

### Physicochemical characteristics and salt tolerance of GXDK6

GXDK6 was incubated in YPD medium containing 0%–18% NaCl for 16–48 h and then observed using an optical microscope and a scanning electron microscope (Carl Zeiss AG, Oberkochen, Germany). The salt removal rate of GXDK6 was also investigated by conductivity method. The results helped determine the most suitable concentration of NaCl to stress GXDK6 for subsequent experiments.

### Genome sequencing analysis of GXDK6

The whole genomic DNA of GXDK6 was extracted using the CTAB method as reported by Van et al. (14), with slight modifications. The purity of the extracted DNA was verified by polymerase chain reaction (PCR) and agarose gel electrophoresis (43). The ITS gene was amplified by PCR using ITS1 (5′-TCCGTAGGTGAACCTGCGG-3′)/ITS4 universal primers (5′-TCCTCCGCTTATTGATATGC-3′) (44). The ITS sequencing data of GXDK6 were deposited to the National Microbiology Data Center database (http://nmdc.cn) under the accession number NMDCN000022O.

Whole-genome sequencing and analysis of GXDK6 were performed by BGI Genomics Co., Ltd (Shenzhen, China). An online software called FastQC (http://www.bioinformatics.babraham.ac.uk/projects/fastqc) was used for the quality control of the second-generation sequencing downtime data. BUSCO software (http://busco.ezlab.org, v3.0.2) was used to complete the sequence comparison of the genome sequences to determine the percentage of single-copy genes in the total single-copy genes. Hmmscan software was used to predict the presence of carbohydrate-active enzyme (http://www.cazy.org) genes in the genome sequence (45, 46). The whole-genome sequencing data of GXDK6 were deposited to the National Microbiology Data Center database (http://nmdc.cn) under accession number NMDC60014229 (14).

### Transcriptome sequencing analysis of GXDK6

When GXDK6 was incubated under 0%–10% NaCl for 16 or 48 h, the strains were centrifuged at 8000× g for 10 min and then transferred into liquid nitrogen for rapid freezing. The total RNA of GXDK6 was extracted using the TRIzol method (47). Subsequently, the extracted RNA was sequenced and analyzed by Guangzhou Gene Denovo Biotechnology Co., Ltd (Guangzhou, China).

### Proteome sequencing analysis of GXDK6

GXDK6 was incubated under 0%, 5%, and 10% NaCl stress for 16 h. The strains were then collected by centrifugation (8000× g), and the proteins of GXDK6 were extracted by SDT lysis methods (48). Proteome sequencing analysis of the extracted proteins was conducted by Guangzhou Gene Denovo Biotechnology Co., Ltd (Guangzhou, China) by using tandem mass tag-based quantitative proteomics (49).

### Metabolome analysis of GXDK6

GXDK6 cells were removed by centrifugation at 12000× g for 10 min after incubation for 16 or 48 h. The zymotic fluid was then filtered by a 0.45 μm microporous membrane (organic system) and placed in a 1.5 mL centrifuge tube. Subsequently, 500 μL of the above zymotic fluid from each condition was transferred into new 1.5 mL centrifuge tubes, and 5 μL of ribitol solution (GC level) at a concentration of 1 mg/mL was added in the tubes as the internal standard. Afterwards, all the samples were concentrated in a freeze centrifugal concentrator until they were completely dried to powder or flocculent, and the samples were used for gas chromatography-mass spectrometry (GC-MS) detection and analysis.

All the samples required a two-step pretreatment before GC-MS detection (50). First, alkylation reaction of the samples was performed by adding 80 μL methoxyamine pyridine hydrochloride solution (a special reagent for GC analysis) with a concentration of 20 mg/mL and then oscillating in a rotary shaker at 200 rpm and 37 °C for 120 min. Second, when the alkylation reaction was completed, 80 μL of MSTFA was added in the samples for derivatization reaction, which was also conducted in a rotary shaker at 200 rpm and 37 °C for 120 min. Subsequently, the samples were centrifuged at 12000× g for 10 min to collect the supernatant for GC-MS detection.

The samples above were detected using DSQ II single quadrupole GC-MS (Thermo Fisher Company). The injection port temperature of the instrument was maintained at 250 °C. One μL of the derivative sample was extracted and injected into the dodecyl benzene sulfonic acid column (length: 30 mm × inner diameter: 250 μm × thickness: 0.25 μm). Under the direct ionization mode of 70 eV ionization energy and 8000 V accelerating voltage, the source temperature of the mass spectrometer was maintained at 250 °C. In the experiments of full scan and selective ion recording, the temperature of the quadrupole was kept at 150 °C. The initial temperature of the gas chromatograph was set at 85 °C for 5 min, and it was then increased to 330 °C at a rate of 15 °C/min. Helium was used as the carrier gas with a constant flow rate of 1 mL/min, the operating range of mass spectrometry was 50–600 M/Z (51).

### Data analysis

Data fitting and mapping analysis were performed using Origin 9.0 software. Statistical analysis of other experimental data was performed using SPSS 17.0, and p value < 0.05 indicated significant differences. For data obtained by GC-MS, Xcalibur software was used to extract the peak area of the total ion chromatogram, and then principal component analysis and independent component analysis were conducted (52). The differential metabolites were analyzed on GraphPad Prism v5.01 (La Jolla, CA, USA), and heatmap analysis was performed on R software.

## SUPPLEMENTAL METERIAL

**Additional file 1**. Differential genes of GXDK6 under NaCl stress.

**Additional file 2.** Differential proteins of GXDK6 under NaCl stress.

**Additional file 3.** Metabolomics analysis of GXDK6 under NaCl stress.

## ACKNOWLEDGMENTS

The authors would like to thank Guangzhou Gene Denovo Biotechnology Co., Ltd. for the help of high throughput sequencing and analysis of omics data.

This work was supported by the National Natural Science Foundation of China (Grant No. 31760437), Science and Technology Basic Resources Investigation Program of China (Grant No. 2017FY100704), Natural Science Fund for Distinguished Young Scholars of Guangxi Zhuang Autonomous Region of China (Grant No. 2019GXNSFFA245011), Natural Science Foundation of Guangxi Zhuang Autonomous Region of China (Grant No. 2018GXNSFAA050090) and the Innovation Project of Guangxi Graduate Education (YCBZ2021012).

The authors declare that they have no competing interests.

## REFERENCES

1. Shaw A, Joe, Lam, Felix H, Hamilton, Maureen, Consiglio, Andrew. 2016. Metabolic engineering of microbial competitive advantage for industrial fermentation processes. Science 353:583–586. https://doi.org/10.1126/science.aaf6159.

2. Wang Z, Chen X, Liu S, Zhang Y, Zhang. 2019. Efficient biosynthesis of anticancer polysaccharide by a mutant *Chaetomium globosum* ALE20 via non-sterilized fermentation. Int J Biological Macromolecules 136:1106–1111. https://doi.org/10.1016/j.ijbiomac.2019.06.186.

3. Seo HB, Kim SS, Lee HY, Jung KH. 2009. High-level production of ethanol during fed-batch ethanol fermentation with a control of aeration rate and nonsterile glucose powder feeding of *Saccharomyces cerevisiae*. Biotech & Biopro Eng 14:591–598. https://doi.org/10.1007/s12257-008-0274-2.

4. Xu DB, Ye WW, Han Y, Deng ZX, Hong K. 2014. Natural products from mangrove actinomycetes. Marine Drugs 12:2590–2613. https://doi.org/10.3390/md12052590

5. He B, Long M, Hu Z, Li H, Zeng B. 2018. Deep sequencing analysis of transcriptomes in *Aspergillus oryzae* in response to salinity stress. Applied Micro & Biotech 102:1–10. https://doi.org/10.1007/s00253-017-8603-z.

6. Akshya S, Sukesh CS. 2017. Physiological basis for the tolerance of yeast *Zygosaccharomyces bisporus* to salt stress. HAYATI J Biosciences 24:176–181. https://doi.org/10.1016/j.hjb.2017.11.001.

7. Matsushika A, Suzuki T, Goshima T, Hoshino T. 2017. Evaluation of *Saccharomyces cerevisiae* GAS1 with respect to its involvement in tolerance to low pH and salt stress. J Biosci & Bioeng 124:164–170. https://doi.org/10.1016/j.jbiosc.2017.03.004.

8. Yang HL, Liao YY, Zhang J, Wang XL. 2019. Comparative transcriptome analysis of salt tolerance mechanism of Meyerozyma *guilliermondii* W2 under NaCl stress. 3 Biotech 9:286. https://doi.org/10.1007/s13205-019-1817-2.

9. Hou L, Wang M, Wang C, Wang C, Wang H. 2013. Analysis of salt-tolerance genes in *Zygosaccharomyces rouxii*. Applied Biochem & Biotech 170:1417–1425. https://doi.org/10.1007/s12010-013-0283-2.

10. Rea SM, Kaksonen A & H. 2015. Salt-tolerant microorganisms potentially useful for bioleaching operations where fresh water is scarce. Minerals Eng 75:126–132. https://doi.org/10.1016/j.mineng.2014.09.011.

11. Eichner J, Rosenbaum L, Wrzodek C, HU Häring, Lehmann R. 2014. Integrated enrichment analysis and pathway-centered visualization of metabolomics, proteomics, transcriptomics, and genomics data by using the InCroMAP software. J Chromatogr B Analyt Technol Biomed Life 966:77–82. https://doi.org/10.1016/j.jchromb.2014.04.030.

12. Peinemann JC, Rhee C, Shin SG, Pleissner D. 2020. Non-sterile fermentation of food waste with indigenous consortium and yeast – Effects on microbial community and product spectrum. Biores Tech 306:123175. https://doi.org/10.1016/j.biortech.2020.123175.

13. Bi J, Liu S, Du G, Chen J. 2015. Bile salt tolerance of *Lactococcus lactis* is enhanced by expression of bile salt hydrolase thereby producing less bile acid in the cells. Biotech Let 38:659–665. https://doi.org/10.1007/s10529-015-2018-7.

14. Mo X, Cai X, Hui Q, Sun H, Jiang C. 2021. Whole genome sequencing and metabolomics analyses reveal the biosynthesis of nerol in a multistress-tolerant *Meyerozyma guilliermondii* GXDK6. Micro Cell Fact 20:1–11. https://doi.org/10.1186/s12934-020-01490-2.

15. Abou-Elela SI, Kamel MM, Fawzy ME. 2010. Biological treatment of saline wastewater using a salt-tolerant microorganism. Desalination 250:1–5. https://doi.org/10.1016/j.desal.2009.03.022.

16. Wang D, Hao Z, Zhao J, Jin Y, Huang J, Zhou R, Wu C. 2019. Comparative physiological and transcriptomic analyses reveal salt tolerance mechanisms of *Zygosaccharomyces rouxii*. Process Biochem 82:59–67. https://doi.org/10.1016/j.procbio.2019.04.009.

17. Wei X, Chi Z, Liu GL, Hu Z, Chi ZM. 2020. The genome-wide mutation shows the importance of cell wall integrity in growth of the psychrophilic yeast *Metschnikowia australis* W7-5 at different temperatures. Microbial Ecology 8: 1–15. https://doi.org/10.1007/s00248-020-01577-8.

18. Zhang A, Kong Q, Cao L, Chen X. 2010. Effect of FPS1 deletion on the fermentation properties of *Saccharomyces cerevisiae*. Let in Appl Micro 44:212–217. https://doi.org/10.1111/j.1472-765X.2006.02041.x.

19. Hohmann S, Krantz M, Nordlander B. 2007. Yeast osmoregulation. Methods in Enzymol 428:29–45. https://doi.org/10.1016/S0076-6879(07)28002-4.

20. Parmar JH, Bhartiya S, Venkatesh KV. 2011. Characterization of the adaptive response and growth upon hyperosmotic shock in *Saccharomyces cerevisiae*. Molecular Biosys 7:1138–1148. https://doi.org/10.1039/c0mb00224k.

21. Wang, Y., Tao, F., Xu, P. 2014. Glycerol dehydrogenase plays a dual role in glycerol metabolism and 2,3-butanediol formation in *Klebsiella pneumoniae*. J Biolog Chem 289:6080. https://doi.org/10.1074/jbc.M113.525535.

22. Mane SP, Evans C, Cooper KL, Crasta OR, Folkerts O, Hutchison SK. 2009. Transcriptome sequencing of the microarray quality control (MAQC) RNA reference samples using next generation sequencing. Bmc Genomics 10: 264–265. https://doi.org/10.1186/1471-2164-10-264.

23. Seo SH, Park SE, Yoo SA, Lee KI, Na CS, Son HS. 2016. Metabolite profiling of *Makgeolli* for the understanding of yeast fermentation characteristics during fermentation and aging. Proc Biochem. 51:1363–1373. https://doi.org/10.1016/j.procbio.2016.08.005.

24. Ness F, Aigle M. 1995. RTM1: a member of a new family of telomeric repeated genes in yeast. Genetics 140:945–956. https://doi.org/10.1101/gad.9.13.1667.

25. Cassanova N, O’Brien KM, Stahl BT, Mcclure T, Poyton RO. 2005. Yeast flavohemoglobin, a nitric oxide oxidoreductase, is located in both the cytosol and the mitochondrial matrix: effects of respiration, anoxia, and the mitochondrial genome on its intracellular level and distribution. J Biological Chem 280:7645–7653. https://doi.org/10.1074/jbc.M411478200.

26. Liu L, Si L, Meng X, Luo L. 2015. Comparative transcriptomic analysis reveals novel genes and regulatory mechanisms of *Tetragenococcus halophilus* in response to salt stress. J Industrial Micro & Biotech 42:601–616. https://doi.org/10.1007/s10295-014-1579-0.

27. Vasim A, Verma MK, Shashank G, Vibha M, Chauhan NS. 2018. Metagenomic profiling of soil microbes to mine salt stress tolerance genes. Front in Micro 9:159–159. https://doi.org/10.3389/fmicb.2018.00159.

28. Lv G, Jiang C, Liang T, Tu Y, He B. 2020. Identification and expression analysis of sugar transporter gene family in *Aspergillus oryzae*. Int J Genomics 2020:1–15. https://doi.org/10.1155/2020/7146701.

29. Aguilera J, Prieto J. 2001. The *saccharomyces cerevisiae* aldose reductase is implied in the metabolism of methylglyoxal in response to stress conditions. Current Genetics 39:273–283. https://doi.org/10.1007/s002940100213.

30. Su H, Feng J, Lv J, Liu Q, Xie S. 2021. Molecular mechanism of lipid accumulation and metabolism of *oleaginous chlorococcum sphacosum* GD from soil under salt stress. Int J Molecular Sciences 22:1304. https://doi.org/10.3390/ijms22031304.

31. Adler L, Blomberg A, Nilsson A. 1985. Glycerol metabolism and osmoregulation in the salt-tolerant yeast *Debaryomyces hansenii*. J Bacteriology 162:300–306. https://doi.org/10.1128/JB.162.1.300-306.

32. Potts MB, Mcmillan EA, Rosales TI, Kim HS, Ou YH, Toombs JE. 2015. Mode of action and pharmacogenomic biomarkers for exceptional responders to Didemnin B. Nat. Chem. Bio 11:401–408. https://doi.org/10.1038/nchembio.1797.

33. Malloy PJ, Fe Ldman D. 2011. The role of vitamin d receptor mutations in the development of alopecia. Molecular & Cellular Endocrinology 347:90–96. https://doi.org/10.1016/j.mce.2011.05.045.

34. Chambers ES, Suwannasaen D, Mann EH, Urry Z, Richards DF, Lertmemongkolchai G. 2014. 1α,25-dihydroxyvitamin D3 in combination with transforming growth factor-β increases the frequency of Foxp3 regulatory T cells through preferential expansion and usage of interleukin-2. Immunol 143:52–60. https://doi.org/10.1111/imm.12289.

35. Henne WA, Kularatne SA, Ayala-López W, Doorneweerd DD, Stinnette TW, Lu Y. 2012. Synthesis and activity of folate conjugated Didemnin B for potential treatment of inflammatory diseases. Bioorganic & Medicinal Chem Let 22: 709–712. https://doi.org/10.1016/j.bmcl.2011.10.042.

36. Jinqi, Luo, Fang, Jiang, Weizhen, Fang. 2016. Optimization of bioconversion conditions for vitamin D3 to 25-hydroxyvitamin D using *Pseudonocardia autotrophica* CGMCC5098. Biocatalysis and Biotransformation. 35:11–18. https://doi.org/10.1080/10242422.2016.1268130.

37. Peinemann JC, Rhee C, Shin SG, Pleissner D. 2020. Non-sterile fermentation of food waste with indigenous consortium and yeast-Effects on microbial community and product spectrum. Biores Tech 306:123175. https://doi.org/10.1016/j.biortech.2020.123175.

38. Na LA, Dong WA, Xlj A, Feng C, Stya C. 2012. Regulation of lipid metabolism in the snow *alga chlamydomonas nivalis* in response to NaCl stress: An integrated analysis by cytomic and lipidomic approaches. Process Biochem 47:1163–1170. https://doi.org/10.1016/j.procbio.2012.04.011.

39. Roy S, Chakraborty U. 2014. Salt tolerance mechanisms in salt tolerant grasses (STGs) and their prospects in cereal crop improvement. Botanical Studies 55:31. https://doi.org/10.1186/1999-3110-55-31.

40. Liu N, Chen AP, Zhong NQ, Wang F, Xia GX. 2010. Functional screening of salt stress-related genes from *Thellungiella halophila* using fission yeast system. Physiologia Plantarum 129:671–678. https://doi.org/10.1111/j.1399-3054.2007.00857.x.

41. Li C, Zhao D, Yan J, Zhang N, Li B. 2021. Metabolomics integrated with transcriptomics: assessing the central metabolism of marine red yeast *Sporobolomyces pararoseus* under salinity stress. Arch Microbiol 203:889–899. https://doi.org/10.1007/s00203-020-02082-9

42. Van B, Schreckhise RW, White TC, Bowden RA, Myerson D. 1998. Comparison of six extraction techniques for isolation of DNA from filamentous fungi. Medical Mycology 36:299–303. https://doi.org/10.1111/j.1365-280X.1998.00161.x.

43. Ehrhardt DW, Frommer WB. 2012. New technologies for 21st century plant science. Plant Cell 24:374–394. https://doi.org/10.1105/tpc.111.093302.

44. Dupont D, Gaucherand P, Wallon M. 2019. Fortuitous diagnosis of *Trichomoniasis* by PCR using panfungal primers. Int J Infectious Diseases 90: 234–236. https://doi.org/10.1016/j.ijid.2019.11.008

45. Lagesen K, Hallin P, Rodland EA, Staerfeldt HH, Rognes T, Ussery DW. 2007. RNAmmer: consistent and rapid annotation of ribosomal RNA genes. Nucl Aci Res 35:3100–3108. https://doi.org/10.1093/nar/gkm160.

46. Lombard V, Golaconda RH, Drula E, Coutinho PM, Henrissat B. The carbohydrate-active enzymes database (CAZy) in 2013. Nucl Aci Res 42:D490–D495. https://doi.org/10.1093/nar/gkt1178.

47. Villa-Rodríguez E, Ibarra-Gámez C, Sergio SV. 2018. Extraction of high-quality RNA from *Bacillus subtilis* with a lysozyme pre-treatment followed by the trizol method. J Microbiological Methods 147:14–16. https://doi.org/10.1016/j.mimet.2018.02.011.

48. Zhu Y, Xu H, Chen H, Xie J, Shi M, Shen B. 2014. Proteomic analysis of solid pseudopapillary tumor of the pancreas reveals dysfunction of the endoplasmic reticulum protein processing pathway. Molecular & Cellular Proteomics 13:2593–2603. https://doi.org/10.1074/mcp.M114.038786.

49. Myers SA, Klaeger S, Satpathy S, Viner R, Carr SA. 2018. Evaluation of advanced precursor determination for tandem mass tag (TMT)-based quantitative proteomics across instrument platforms. J Proteome Res 18:542–547. https://doi.org/10.1021/acs.jproteome.8b00611.

50. Zhe Wang, Min-Yi Li, Zhi-Xue Peng. 2016. GC-MS-based metabolome and metabolite regulation in serum-resistant *streptococcus agalactiae*. J Proteome Res 1:22–46. https://doi.org/10.1021/acs.jproteome.6b00215.

51. Li H, Huang X, Zeng Z, Peng XX, Peng B. 2016. Identification of the interactome between fish plasma proteins and *Edwardsiella tarda* reveals tissue-specific strategies against bacterial infection. In J Biochem & Cell Biol 78: 260–267. https://doi.org/10.1016/j.biocel.2016.07.021.

52. Yang J, Cheng Q. 2015. A comparative study of independent component analysis with principal component analysis in geological objects identification, Part I: Simulations. J Geochemical Exploration 149:127–135. https://doi.org/10.1016/j.gexplo.2014.11.014.

